# Supporting-cell vs. hair-cell survival in the human cochlea: Implications for regenerative therapies

**DOI:** 10.1101/2023.04.24.538119

**Authors:** Charanjeet Kaur, McKayla Van Orden, Jennifer T. O’Malley, Pei-zhe Wu, M. Charles Liberman

**Affiliations:** Eaton-Peabody Laboratories, Massachusetts Eye and Ear, Boston, MA 02114; Otopathology Laboratory, Massachusetts Eye and Ear, Boston, MA 02114; Dept of Otolaryngology-Head & Neck Surgery, Harvard Medical School, Boston, MA 02115; Northeastern University, Boston, MA 02115

**Keywords:** Hearing loss, temporal bones, audiograms

## Abstract

Animal studies have shown that the supporting-cells surviving in the organ of Corti after cochlear insult can be transdifferentiated into hair cells as a treatment for sensorineural hearing loss. Clinical trials of small-molecule therapeutics have been undertaken, but little is known about how to predict the pattern and degree of supporting-cell survival based on audiogram, hearing loss etiology or any other metric obtainable pre-mortem. To address this, we systematically assessed supporting-cell and hair cell survival, as a function of cochlear location in 274 temporal bone cases from the archives at the Massachusetts Eye and Ear and compared the histopathology with the audiograms and hearing-loss etiologies. Results showed that supporting-cell survival was always significantly greater in the apical half than the basal half of the cochlea, that inner pillars were more robust than outer pillars or Deiters’ cells, and that total replacement of all supporting cells with a flat epithelium was rare outside of the extreme basal 20% of the cochlea. Supporting cell survival in the basal half of the cochlea was better correlated with the slope of the audiogram than with the mean high-frequency threshold per se: i.e. survival was better with flatter audiograms than with steeply down-sloping audiograms. Cochlear regions with extensive hair cell loss and exceptional supporting cell survival were most common in cases with hearing loss due to ototoxic drugs. Such cases also tended to have less pathology in other functionally critical structures, i.e. spiral ganglion neurons and the stria vascularis.

**Highlights:**

- Supporting cell survival was systematically assessed in 274 human cochleas
- Supporting cell survival was better with flat than with down-sloping audiograms
- Supporting cell survival was most robust when hearing loss was from ototoxic drugs
- Ototoxic cases also showed less pathology in other critical cochlear structures
- The data can inform clinical trials for regeneration via supporting cell conversion

## 1. Introduction

Much of the threshold elevations that define permanent sensorineural hearing loss in humans are due to irreversible loss of, or damage to, the hair cells (Bredberg 1968, Merchant and Nadol 2010, Wu, O’Malley et al. 2020), which are responsible for transducing sound-evoked vibrations of the sensory epithelium into electrical activity for transmission to the central nervous system. Hair cells are vulnerable to damage and/or degeneration from a number of cochlear insults. These include: ototoxic drugs such as aminoglycoside antibiotics (Dallos and Harris 1978, Johnsson, Hawkins et al. 1981) or platinum-containing anti-cancer medications (Taudy, Syka et al. 1992) or overexposure to loud sounds (Johnsson and Hawkins 1976, McGill and Schuknecht 1976). Hair cell death also underlies much of the threshold elevations of age-related hearing loss (Schuknecht and Gacek 1993, Wu, O’Malley et al. 2020).

In general, hair cells are among the most vulnerable cells in the inner ear. Thus, there can be fractional or total hair cell loss in regions of the sensory epithelium where normal or near-normal architecture of the organ of Corti’s supporting cells remains (Liberman and Kiang 1978, Suzuka and Schuknecht 1988, Sugawara, Corfas et al. 2005). The issue of supporting-cell survival in sensorineural hearing loss has taken on added significance since the discovery of spontaneous, post-traumatic hair-cell regeneration in birds, because that regeneration arises from cell-division and/or trans-differentiation of surviving supporting cells in the sensory epithelium of the hearing organ (Warchol and Corwin 1996). Those discoveries in non-mammalian ears, in turn, have led to the demonstration in adult mice that, after noise-induced hair cell destruction, forced trans-differentiation of surviving supporting cells into hair-cell like cells can be elicited by local cochlear delivery of a small-molecule therapeutic, and that the modest numbers of trans-differentiated hair cells can be associated with a modest improvement in cochlear threshold sensitivity (Mizutari, Fujioka et al. 2013, McLean, Yin et al. 2017). Most importantly, these preclinical studies have also spawned development of human therapeutics for sensorineural hearing loss, and one such drug has even moved into clinical trials (McLean, Hinton et al. 2021). However, the clinical trials to date have not been highly successful.

The general resistance of the organ of Corti’s supporting cells to cochlear insults in cases of sensorineural hearing loss has been documented in both animal and human otopathology (Suzuka and Schuknecht 1988, Raphael and Altschuler 1991, Sugawara, Corfas et al. 2005, Kamakura, O’Malley et al. 2018, deTorres, Olszewski et al. 2019), but it has never been systematically analyzed in large numbers of cases. Here, we set out to correct that knowledge gap by assessing supporting cell and hair cell survival in a large number of cases (n=274) of sensorineural hearing loss from the archive of human inner ear specimens in the Temporal Bone Library at the Massachusetts Eye and Ear. (Merchant, McKenna et al. 2008). We included cases with a variety of hearing loss etiologies, and with a variety of audiometric patterns, so that we could assess whether the probability of seeing exceptional supporting cell survival in the face of significant hair cell loss was dependent on any criteria that could be assessed pre-mortem and factored into the inclusion criteria for future clinical trials.

Results showed 1) that complete loss of all recognizable supporting-cell structures is rare in all but the most profound cases of hearing loss, 2) that supporting cell loss is minimal in the apical half of the cochlea regardless of audiometric pattern, and 3) that supporting cell loss in the basal half of the cochlea is better predicted by considering the slope of the audiogram, i.e. the difference between high-frequency and low-frequency pure-tone averages, than by considering the high-frequency thresholds in isolation. Comparison of data from different hearing loss etiologies suggests that patients with ototoxic-induced hearing loss might be better candidates than those with noise-induced damage.

## 2. Methods

### 2.1. Case selection

We studied 274 human temporal bones (142 male ears and 132 female ears) from individuals aged 10 to 104 yrs, selected from the archives at the Massachusetts Eye and Ear (MEE). To be included in the study, the histological preservation had to be adequate to reliably assess the condition of hair cells and supporting cells, and the associated audiogram had to be one of three general configurations: 1) “descending”, with thresholds better than 40 dB at low frequencies (0.25, 0.5 and 1 kHz) and worse than 50 dB at high frequencies (2, 4 and 8 kHz); 2) “flat”, with a threshold standard deviation across all frequencies < 10, or 3) “profound”, with all thresholds worse than 90 dB. The first audiometric pattern was similar to that used as an inclusion criterion in the human clinical trial (McLean, Hinton et al. 2021). In some analyses, we further limited the sample to those for which the audiogram was obtained with 6 yrs of death, and to those for which the air-bone gap was < 15 dB, averaged across all frequencies (see Figure Captions). All procedures and protocols for the study of archived human tissue were approved by the Institutional Review Board of the Massachusetts Eye and Ear.

### 2.2. Histological analysis

Each case consists of a set of every 10^th^ section (20 µm thickness) through the decalcified and celloidin-embedded temporal bone, stained with hematoxylin and eosin. The cochlea in each case was graphically reconstructed (Merchant and Nadol, 2010) to compute the percent distance along the spiral of each section through the cochlear duct, which was then converted into frequency using a cochlear map for human (Greenwood, 1990), modified to produce an apical-most best frequencies of 100 rather than 20 Hz. This modification was introduced because 1) the lowest best frequencies ever recorded from the mammalian auditory nerve (including macaque (Shera, Bergevin et al. 2011), cat (Liberman 1978), gerbil (Schmiedt 1989), guinea pig (Tsuji and Liberman 1997) and chinchilla (Henry, Kale et al. 2016)) are all > 100 Hz; and 2) if human auditory-nerve fibers are comparable, those tuned to 100 Hz would respond to 20 Hz tones at SPLs appropriate to the human audiogram.

Supporting-cell survival was rated semi-quantitatively, by observers blinded to all associated case data, according to a 5-point rating scale described in more detail in Results. Supporting cells were assessed in all sections through the organ of Corti, from all slides in the set, i.e. at ∼ 100 points along the cochlear spiral. Analysis was performed using digital images scanned by an Aperio AT2 (Leica) using a planapo 20x objective (N.A. 0.8). Fractional survival of Inner and outer hair cells was also assessed at all cochlear locations in most cases, using the original microscope slides and imaged with a 100X immersion objective (N.A. 1.3) and differential interference contrast on a Nikon E800 microscope. A detailed description of the criteria used in these hair cell counts is available in a prior publication (Wu, Wen et al. 2019). In a subset of the cases, i.e. those with poor hair cell survival yet excellent supporting-cell survival, the condition of several other key structures of the cochlear duct was evaluated semi-quantitatively according to a rating scale described more fully in Figure Captions. For this semi-quantitative analysis, in each case, we examined one mid-modiolar section (which includes five views of the organ of Corti at loci from 40% to the 100% of the distance from the base), and a second section roughly halfway between the mid-modiolar and the ventral-most section (which includes views at roughly the 12.5 and 30% loci). At each locus, an experienced observer estimated the survival of each structure on a 3-point scale: 0 - 33%, 33 - 66% or 66 - 100%.

### 2.3. Statistical analysis

K-mean cluster analysis was performed on the binned supporting cell data using the *kmeans* function in Matlab. Regression analyses and t-tests were performed in Excel, and two-way ANOVAs were carried out in GraphPad Prism using a mixed-effects model with Gasser-Greenhouse correction.

## 3 Results

### 3.1. The rating scale for assessment of supporting cell condition

In archival temporal bone material stained with hematoxylin and eosin, a large amount of cellular detail is extractable using high-power oil-immersion lenses and differential interference contrast microscopy. Fractional hair cell survival can be accurately assessed in these semi-serial 20 µm sections by optical sectioning with high-N.A. objectives (Wu, Wen et al. 2019). This highly quantitative approach could be applied to inner and outer pillar cells and the Deiters cells, which are recognizable by virtue of the highly eosinophilic microtubular bundles that form a sort of endoskeleton within each of these cell types (Slepecky, Henderson et al. 1995) (Fig. 1). However, such a meticulous quantification of the fractional survival of these supporting cells was prohibitively time consuming for a study that aimed to provide a broad overview by surveying a large number of cases. Thus, we devised a semi-quantitative, 5-point rating scale with which all sections from each case could be assessed in ∼ 1 hour. Early in the process, we compared ratings assigned by two users until the inter-observer and inter-trial reproducibility was acceptable (r^2^ = 0.88 for ratings on the same slides from two observers).

**Figure 1:**
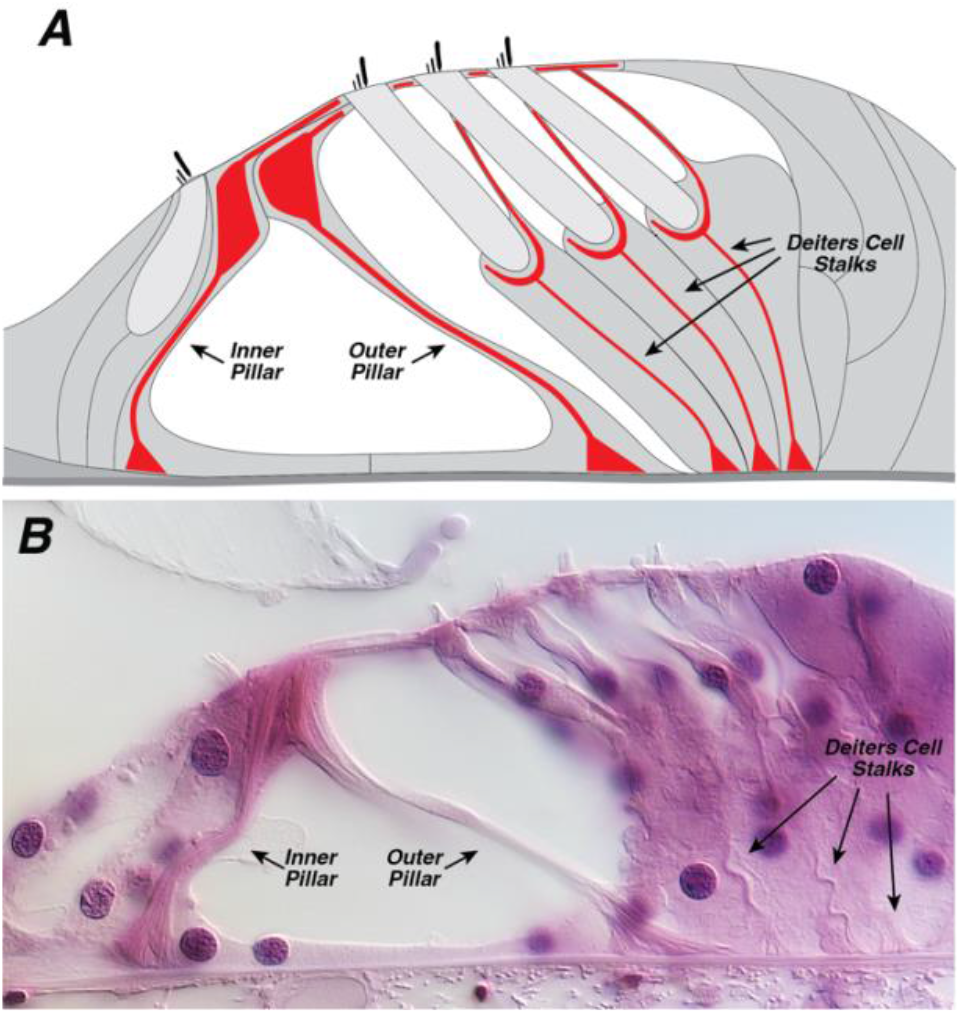
The supporting cells that are the focus of the present study. **A**: Schematic of the organ of Corti, highlighting (in red) the microtubular clusters in pillar and Deiters cells. **B**: Photomicrograph of the human organ of Corti from the upper basal turn, showing how these microtubular clusters appear in celloidin sections stained for hematoxylin and eosin and examined with differential interference contrast microscopy.

As illustrated in Figure 2, we defined the five-point rating scale as follows. A score of 0 was assigned when there was only a low, flat and completely undifferentiated cell layer on top of the basilar membrane. A score of 1 was assigned if there was a mound of epithelial cells present, but no sign of differentiation into supporting cells, i.e. no microtubular bundles indicative of cells which presumably once were pillar or Deiters cells, depending on their position on the basilar membrane. A score of 1.5 was given if the section through the organ of Corti showed epithelial cells present with microtubular clusters suggestive of supporting cells, but no normal architecture, e.g. a completely collapsed tunnel of Corti. As score of 2.0 indicated that the cytoarchitecture was recognizable with a tunnel of Corti, clearly separating outer and inner hair cell regions, but < 50% of supporting cells remained, and a score of 3.0 was assigned if there was normal cytoarchitecture with > 50% of supporting cells remaining.

**Figure 2:**
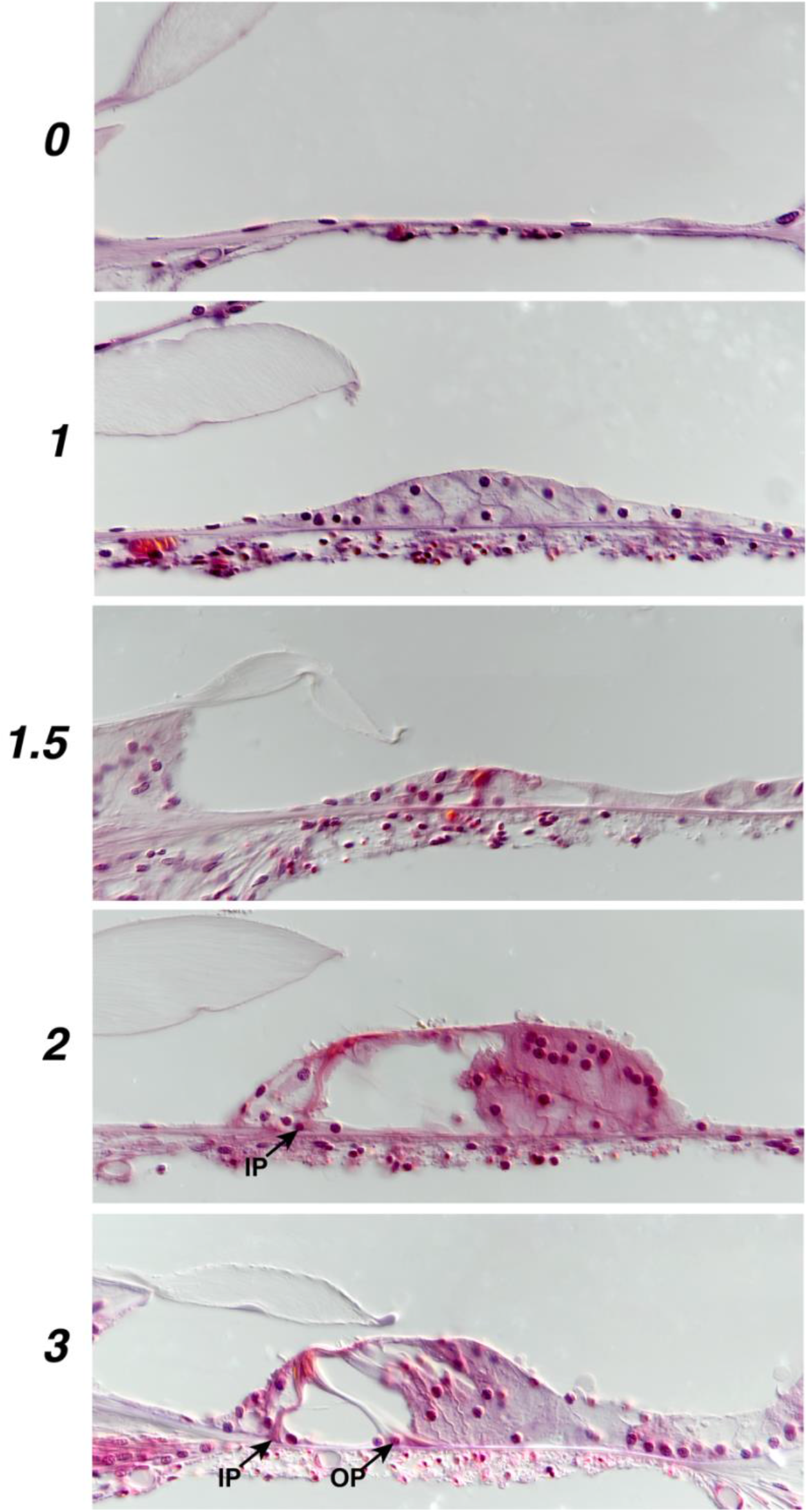
Photomicrographs illustrating the condition of the supporting cells corresponding to the 5-point rating scale used. See text for further details. IP and OP stand for inner pillar and outer pillar, respectively.

For each section receiving a 2.0, the observer noted which of the three supporting cell types dominated the loss. Across all the cases analyzed, there was a clear progression: Deiters cells were most often selectively missing (68% of the time), outer pillar cells were selectively missing only 32% of the time, and inner pillars were never missing unless at least one of the other two types had also disappeared. The exemplar of a “2” rating in Figure 2 show an example in which several of the outer pillars are missing, while the inner pillars remain.

### 3.2. Analysis by K-means Clustering

One way to gain an overview of the patterns of supporting-cell degeneration is to plot the data in each case as a function of cochlear location and use k-means clustering to divide the cases into groups based on the similarity of their histopathological patterns. The results of the cluster analysis (Fig. 3A,B) show several overarching trends. First, supporting-cell survival is almost always better in the apical (low-frequency) half of the cochlea than in the base. Indeed, in virtually all the cases, there is at least some remnant of supporting-cell structure (ratings > 1.5) everywhere in the apical half, and on average the apical organ of Corti retains a normal architecture with more than half of the supporting cells remaining (Fig 3B). Second, replacement of the organ of Corti with a completely undifferentiated epithelium (score of 0) is rarely seen outside of the extreme basal tip of the cochlea. There is sometimes a second focus of severe supporting-cell degeneration in the extreme apex (especially common in cluster 1), which mirrors the additional focus of hair cell degeneration (especially outer hair cells) in the extreme apex (Figs. 3D,E).

**Figure 3:**
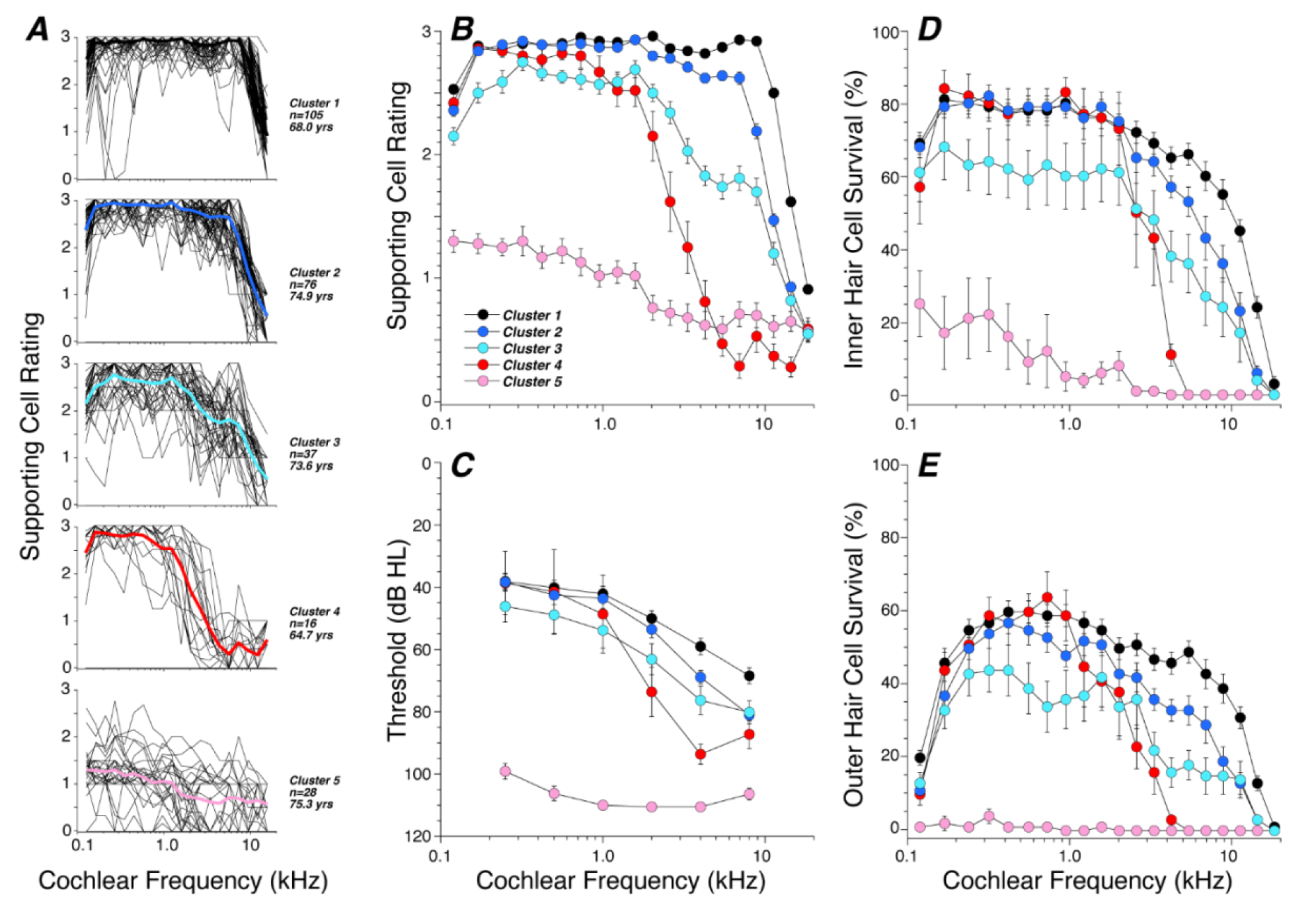
Cluster analysis reveals the overall patterns of supporting-cell survival as a function of cochlear location for all 274 cases in the present study. **A**: Results of K-means clustering. In each case, supporting cell ratings were average into bins of 5% cochlear length (thin black lines), and the mean of each cluster is shown, color coded to match the key in **B. B**: Mean data from **A** are superimposed for ease of comparison, with standard errors added. **C**: Mean audiometric data are shown for each cluster, filtered to include only those with air-bone gap < 15 dB and with test-death interval < 6 yrs. **D**,**E**: Mean IHC and OHC survivals for each cluster.

Not surprisingly, the patterns of supporting-cell degeneration are mirrored in the patterns of audiometric threshold and hair cell loss: i.e. ordering clusters by degree of supporting-cell loss in the basal cochlea is similar to the order of severity in mean high-frequency threshold shifts for the clusters (Fig. 3C) and inner or outer hair cell loss in the basal half of the cochlea (Figs. 3D,E). On the other hand, the order of cluster severity is not obviously mirrored in mean age of the subjects (See keys in Fig. 3A).

### 3.3. Analysis by Audiometric Pattern

Because audiometric data are available pre-mortem as an inclusion criterion in a clinical trial, it is useful to classify cases based on the audiometric pattern. To this end, we devised a simple classification scheme based on the low-frequency pure-tone average (PTA) and the standard deviation of the thresholds across all audiometric frequencies (Fig. 4A). Since “upsloping” audiograms are rare, this strategy divides most audiograms into six non-overlapping groups. By binning the low-frequency PTA (PTA_L_) by 25 dB steps (PTA_L_ ≥ 25, 25 < PTA_L_ ≥ 50, 50< PTA_L_ ≥ 75 and PTA_L_ > 75) and by separating the standard-deviation axis into two groups (SD < 16 vs. SD > 16), we separate the audiograms in the present study into *Normal, Sensory, Descending, Flat Moderate, Flat Severe* and *(Flat) Profound* groups (Fig. 4A,B).

**Figure 4:**
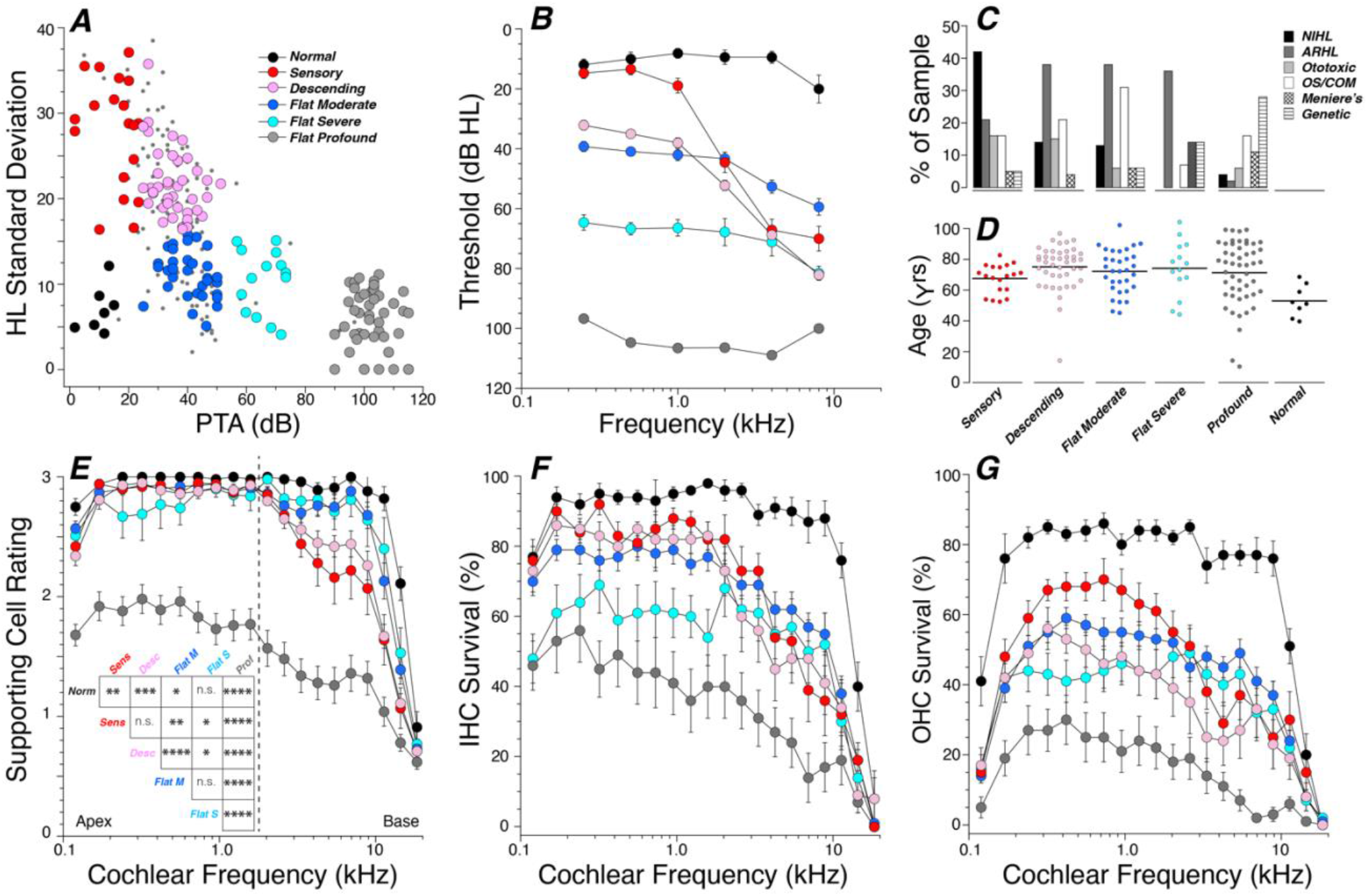
Analysis of supporting-cell survival as a function of audiometric pattern. **A**: Our strategy for audiometric classification separates the groups based on the PTA (average of thresholds at 0.25, 0.5 and 1.0 kHz) and the standard deviation of thresholds across all audiometric frequencies. In each group, cases are removed from subsequent analysis if the air bone gap was > 15 dB or the test-death interval was > 6 yrs (small grey circles). **B**: Mean audiometric thresholds (± SEMs) for each group, after the filtering described for **A. C**,**D**: Distribution of etiologies and ages, respectively, for each of the audiometric groups with hearing loss. OS/COM stands for otosclerosis or chronic otitis media. **E**,**F**,**G**: Mean cell rating or survival percentages (± SEMs) for supporting cells, IHCs, or OHCs, respectively, averaged into bins corresponding to 5% of cochlear length. Matrix inset to **E** shows the significance of the pairwise intergroup differences in supporting cell ratings for the basal half of the cochlea, i.e. all points to the right of the dashed line: * p < 0.05, ** p < 0.005, *** p < 0.0005, **** p < 0.0001.

The *Sensory* group is similar to that described by the Schuknecht and Dubno groups (Pauler, Schuknecht et al. 1988, Dubno, Eckert et al. 2013), except the former provided no explicit numerical boundaries, and the latter’s definition is too restrictive to include many of the audiograms in the present study (Dubno, Eckert et al. 2013). Similarly, our *Flat Moderate* group is similar to Dubno’s *Metabolic* group, but relatively few of our audiograms fit into their precise templates. Our *Descending* group has no obvious counterpart in either of the prior schemes. The dot plots (Fig. 4D) suggest no major differences in ages represented in the different audiometric groups with hearing loss, as borne out by inter-group t-tests (p > 0.05 for all intergroup differences except *Descending* vs. *Sensory*, for which p = 0.04). Not surprisingly, the *Normal* group tends to be younger than all the others.

In the apical half of the cochlea (left of the dashed line in Fig. 4E), although there is generally little loss of supporting-cell architecture, the supporting-cell degeneration is clearly worst in the *Profound* hearing loss group (Fig. 4B; p < 0.005 for all intergroup comparisons by 2-way ANOVA). The same holds for the patterns of inner and outer hair cell loss (Figs. 4D,E), i.e. the mean loss generally increases as the mean threshold shift increases.

In the basal half of the cochlea, where the supporting-cell damage is more severe, the progression of supporting-cell damage deviates from the hearing-loss progression in an interesting way. The supporting-cell damage patterns in the two “flat” audiometric patterns (*Flat Moderate* and *Flat Severe*) are statistically indistinguishable (see matrix inset to Fig. 4E), though the differences in their audiometric patterns are highly significant (Fig. 4B; p < 0.0001). Furthermore, both these “flat” groups show significantly less supporting-cell damage than the two “down-sloping” audiometric groups (*Sensory* and *Descending*), even though both *Sensory* and *Descending* groups have worse high-frequency thresholds than the *Flat Moderate* group.

The notion that the supporting cell survival in the basal half of the cochlea might be better predicted by a measure of audiometric slope rather than the absolute hearing level at high frequencies is tested statistically in Figure 5, where the mean supporting-cell ratings for all ears are regressed against either the mean high-frequency PTA (Fig. 5A) or the difference between high- and low-frequency PTAs (Fig. 5B). Although both correlations are highly significant (p <<< 0.001), the r^2^ value is almost twice as high when supporting-cell survival is regressed against the slope-related metric than when regressed against high-frequency PTA *per se*. Although the correlation between mean supporting-cell damage and age is not significant (Fig. 5C; p = 0.33), there is a significant correlation between supporting-cell damage and duration of deafness (p = 0.002; Figs. 5D).

**Figure 5:**
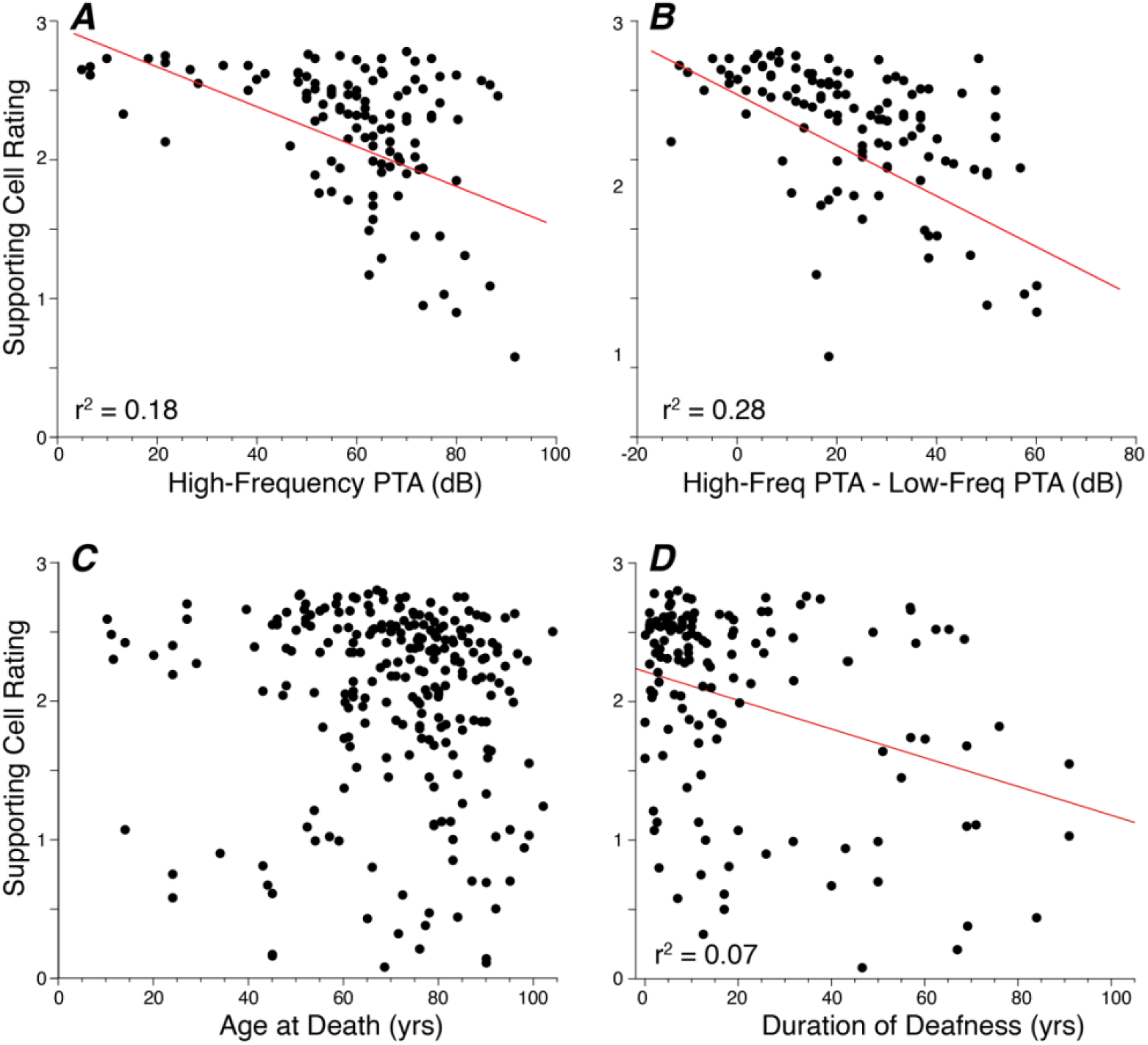
Supporting-cell survival in the basal half of the cochlea is plotted as a function of audiometric thresholds (**A**,**B**), age (**C**) or duration of deafness (**D**). **A**,**B**: Each point represents a different case with air-bone gap < 15 dB and audiogram-death interval < 6 yrs. In **A**, the x-axis is the PTA for 2.0, 4.0 and 8.0, while in **B**, the x axis is the difference between the PTA for 2.0, 4.0 and 8.0 kHz and the PTA for 0.25, 0.5 and 1.0 kHz. **C**,**D**: The y-axis values are as described above, but these panel include cases regardless of audiogram-death interval or air bone gap. Deafness duration (**D**) is based largely on self-report and is unavailable for many of the cases in panel **C**. The best-fit straight line and r^2^ value are shown for those correlations that are statistically significant.

A consideration of how hearing-loss etiology interacts with audiometric shape in our sample of cases is summarized in Figure 4C. Among the four hearing-loss groups for which conductive hearing losses can be effectively screened out by assessment of air-bone gaps (all except *Profound*), the only notable trend is that the *Sensory* group is dominated by cases with a history of noise-induced hearing loss (NIHL). Note that the non-*Profound* SNHL cases with etiologies of Otosclerosis or Chronic Otitis Media (COM) (unfilled bars in Fig. 4C) were able to pass our air-bone gap criterion (< 15 dB mean), because the stapes fixation had been addressed surgically and/or the conductive component of the chronic otitis media (COM) had largely been resolved.

### 3.4. Characterizing cases with exceptional supporting cell survival

Although the overall patterns of supporting-cell survival can be described with mean data, organized either by audiometric groupings (Fig. 4) or cluster analysis (Fig. 3), it is also important to consider the data on a case-by-case basis, to identify those cases with exceptional supporting-cell survival despite considerable hair cell loss. Such cases would presumably be the best candidates for a hair-cell regenerative therapy based on trans-differentiation of supporting cells.

Each point in Figure 6A and 6B compares the mean supporting-cell survival in either the basal or apical half of the cochlea in one case to the mean loss of inner or outer hair cells, respectively, in the same regions of the same cases. There are no points in the upper left half of the plot, because hair cells do not survive in the adult ear in the absence of supporting cells (Mellado Lagarde, Wan et al. 2014). Although most of the points fall near a diagonal, there are a number of points from both the apex and base that fall significantly below that line. To more carefully study cases that might make the best candidates for hair-cell regeneration therapy, we identified those cochlear regions in which the mean supporting-cell rating was > 2.0, while either the outer or inner hair cell survival was < 33%, i.e. those points within the dashed red boxes in Figure 6A and B. As shown in Figure 6C, these inclusion criteria yielded 39 cases with < 33% outer hair cell survival in the base and 18 in the apex, along with 13 case with < 33% inner hair cell survival in the base and 9 cases in the apex. The etiology analysis of panel C suggests that sensorineural hearing loss cases with an ototoxic etiology have the highest probability of presenting with excellent supporting cell survival. Indeed, 12/18 ototoxic cases fell within these inclusion criteria for outer hair cells in the basal turn, arguably the most relevant targets for any regenerative therapies.

**Figure 6:**
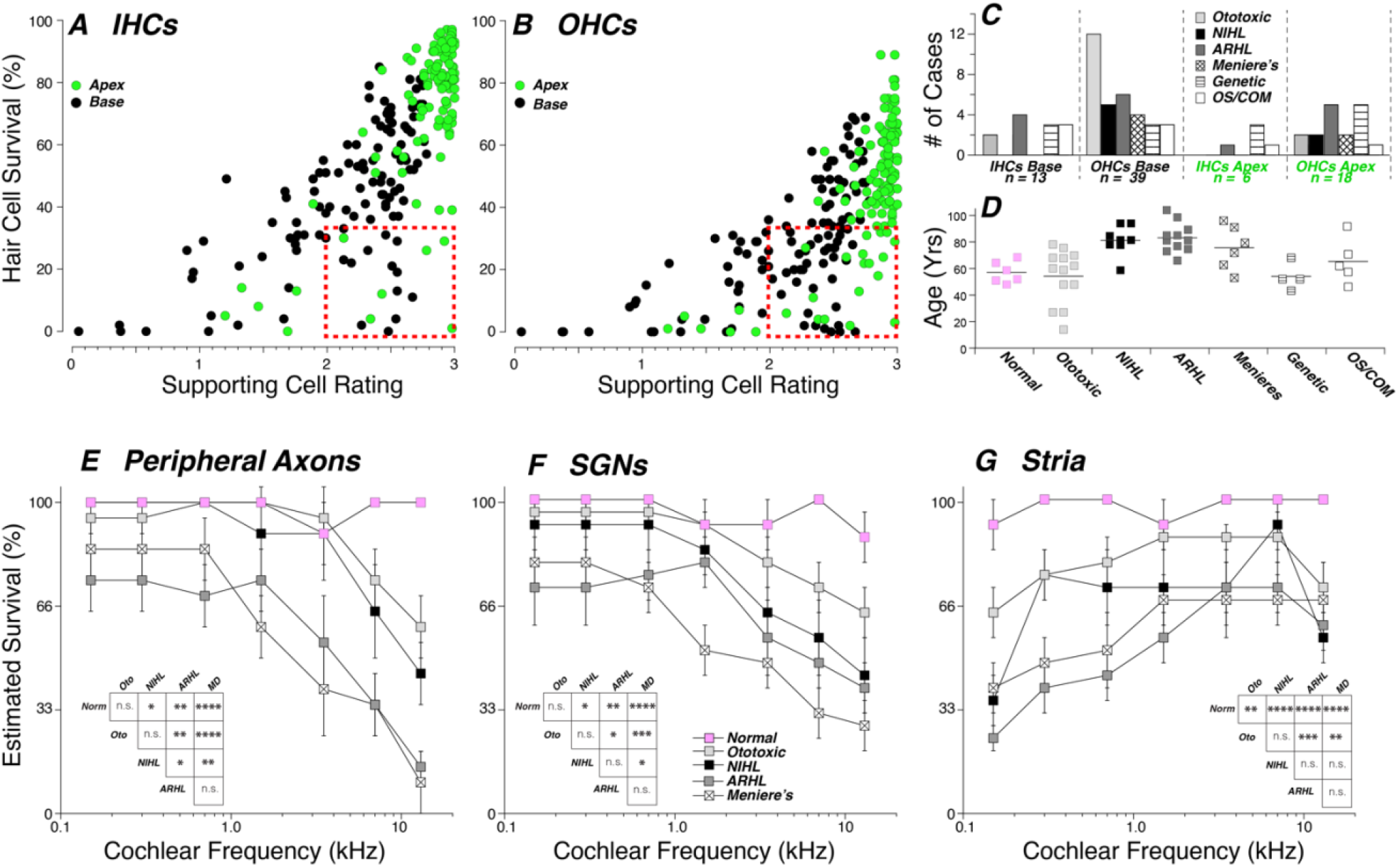
Supporting-cell survival generally follows hair-cell survival, but outlier regions with robust supporting cells despite massive hair cell loss (red boxes in panels **A** and **B**) are interesting candidates for regenerative therapies. **A**,**B**: Each point plots the average SC rating in either the apical or basal half of the cochlea (green or black points) vs. the corresponding mean hair cell survival for either IHCs **(A**) or OHCs (**B**). **C**: Distribution of etiologies for all the candidate points (n = 76 points from 62 cases) enclosed within the red boxes, segregated by hair cell type and cochlear location as indicated. **D**: Ages for all 62 cases yielding points within the red boxes, segregated by etiologies. **E**,**F**,**G**: Mean estimated survival (±SEMs) for peripheral axons, SGNs and the stria vascularis, respectively, for all the cases from each of the four major etiologies represented by the points identified in panels **A**,**B**,**C** and **D**, as compared to the *Normal* group. Matrix insets in **E**,**F**,**G** display the significance of the pairwise intergroup differences as computed via 2-way ANOVA: * p < 0.05, ** p < 0.005, *** p < 0.0005, **** p < 0.0001.

A further complication in the context of hair cell regeneration as a therapy for sensorineural hearing loss is the possible presence of other pathologies among the key cell types in the inner ear. To provide some insight into this, we further analyzed all the candidate cases identified in Figure 6A and B, by performing a semi-quantitative analysis of the condition/survival of auditory-nerve peripheral axons (Fig. 6E), spiral ganglion neurons (SGNs; Fig. 6F), and the stria vascularis (Fig. 6G). Many of these cases showed significant neural degeneration, especially in the basal half of the cochlea, whether with respect to peripheral axons or SGNs; however, the order of increasing severity was ototoxic < NIHL < ARHL < Ménières. The differences between ototoxic and each of the latter two groups was statistically significant for both the peripheral axon and SGN analyses (see matrix inset to Figs. 6E,F). Strial degeneration was prominent in the apical half of the cochlea in many ears (Fig. 6G). Here, the order of increasing severity was ototoxic < NIHL < Ménières < ARHL, but, again, the ototoxic group showed significantly better survival than either ARHL or Ménières groups (see matric inset to Fig. 6G). There was endolymphatic hydrops in all the Ménière’s cases, and occasional collapse of the endolymphatic spaces in a few of the non-Ménière’s cases. In a few cases (10/60 in this group), the tectorial membrane at the basalmost observation point was absent or retracted into the inner sulcus (data not shown).

## 4. Discussion

### 4.1. The importance of supporting cells in hair cell regeneration

The sensory cells in the organ of Corti are surrounded by several classes of supporting cells, many of which span the epithelium from the basilar membrane to the endolymphatic surface (Merchant and Nadol 2010). The supporting cells in direct contact with inner hair cells, i.e. the inner border and inner phalangeal cells, are less highly organized, less well characterized anatomically and less easily recognized than those in the direct contact with the outer hair cells, i.e. inner and outer pillar cells and Deiters cells. The pillar cells and Deiters cells, or their remnants, are easily recognized, even in highly pathological material, due to the characteristic bundle of tightly packed microtubules that runs vertically through each of these cells (Slepecky, Henderson et al. 1995), also essentially spanning the epithelium from basilar membrane to reticular lamina (Fig. 1).

These microtubule-containing supporting-cell types are also especially relevant to ongoing translational studies aiming to trans-differentiate surviving supporting cells from deafened ears into stereocilia-bearing chimeric cells capable of mechano-electric transduction (Mizutari, Fujioka et al. 2013). This is the case, because it is the inner pillar and third-row Deiters cells, in particular, that express Lgr5, the G-protein coupled receptor that is associated with the stem-like properties that predispose them to trans-differentiation when stimulated by the right combination of molecular signals (Mizutari, Fujioka et al. 2013, Bramhall, Shi et al. 2014). For all these reasons, in the present study we concentrated our analysis on the survival of pillar cells and Deiters cells.

The success of preclinical experiments on hair cell regeneration in mice deafened by acoustic trauma (Mizutari, Fujioka et al. 2013) has already inspired a clinical trial using similar molecular strategies to improve hearing in human patients with sensorineural hearing loss (McLean, Hinton et al. 2021). The inclusion criteria for this trial included a stable hearing loss with mean thresholds at 500 - 4000 Hz, no worse than 75 dB, due to either noise exposure or sudden sensorineural hearing loss. For comparison purposes, of the 31 cases in our study with a noise history, all of them met this threshold criterion. Of the 12 cases in the present study with sudden sensorineural hearing loss, only 3 met this threshold criterion, and all were from the flat severe audiometric group. The results reported to date from this clinical trial included a relatively small subject pool (n=15), and showed no significant effect on audiometric thresholds, but a small improvement in the word-recognition scores. This small effect is consistent with the creation of new inner-hair cell like structures, given that inner hair cell loss has little effect on pure-tone thresholds (Lobarinas, Salvi et al. 2013), but should improve discrimination of more complex sounds like speech, assuming that the new hair cells were innervated (Resnik and Polley 2021).

The inclusion criteria in the ongoing trials were derived without any systematic human data on the relation between supporting cell survival and audiometric thresholds or etiology. It was this lack of data that inspired the present study in hopes of clarifying which types of patients might be most suitable for future trials of these and other regenerative therapies, at least with respect to maximizing the probability of significant supporting-cell survival.

### 4.2. Comparison to prior work in animals and humans

There are numerous animal and human studies that comment on supporting cell survival in the course of studies concentrating more on the correlations between hair cell and spiral ganglion neuron survival, e.g. (Suzuka and Schuknecht 1988, Linthicum and Fayad 2009, Rask-Andersen, Liu et al. 2010); however, little systematic analysis has been carried out. Most of that animal work has been done on ototoxic drugs, i.e. aminoglycoside antibiotics or platinum chemotherapeutics in either cats (Leake and Hradek 1988, Xu, Shepherd et al. 1993, Sugawara, Corfas et al. 2005), chinchillas (Sugawara, Corfas et al. 2005), guinea pigs (Forge 1985, Raphael and Altschuler 1991), or mice (Oesterle and Campbell 2009, Taylor, Jagger et al. 2012, Smith-Cortinez, Yadak et al. 2021). Prior human studies have included cases with ototoxic drugs or acoustic trauma (Suzuka and Schuknecht 1988), as well as after cochlear implantation (Kamakura, O’Malley et al. 2018, deTorres, Olszewski et al. 2019), but the human work tends to consider a set of cases, each in isolation, with little attempt to synthesize data to reveal general patterns, and most human otopathological studies ignore the supporting cells (e.g. (Merchant and Nadol 2010)).

Although explicit quantification is lacking, there is qualitative agreement across these human and animal studies that apical regions are more resistant to supporting cell loss than basal regions. Although relatively few animal studies examine survival times more than a few months, at least two reports have studied cases up to five years after an ototoxic drug treatment that killed all the hair cells. Both suggest that duration of deafness is key to supporting-cell survival (Leake and Hradek 1988, Sugawara, Corfas et al. 2005). Thus, for example, an ear at 1 yr post deafening showed supporting-cell death confined to the basal third of the cochlea whereas one at 5 yrs post treatment showed virtually no remaining supporting cells anywhere in the cochlea (Sugawara, Corfas et al. 2005). In the present study, we also saw a significant correlation between supporting-cell survival and duration of deafness (Fig. 5D), however the fraction of the variance accounted for is very small, presumably because there are many different etiologies, degrees of hair cell loss and other uncontrolled variables among these cases.

A few studies have tried to assess cellular health by looking at gene expression via immunohistochemistry in supporting cells surviving after trauma. What evidence there is from animal studies suggests that, if supporting cells remain with a recognizable morphology, they also express key supporting cell markers expressed in normal ears (Oesterle and Campbell 2009, Taylor, Jagger et al. 2012) - including Lgr5 (Smith-Cortinez, Yadak et al. 2021). One human study used immunohistochemistry to assess the health of hair cells and supporting cells surviving in ears with a unilateral CI. As in the animal studies, they concluded that the implantation did not greatly affect the health of the surviving supporting cells, at least as seen by the expression of tubulin (Kamakura, O’Malley et al. 2018).

### 4.3. Insights from the present study

Overall, the present study found that total loss of supporting-cell architecture was relatively rare outside of the extreme basal tip of the cochlea, tuned to frequencies higher than those tested audiometrically (Fig. 3A). In general, the degree of supporting-cell loss mirrored the degree of inner hair loss (Fig. 3), meaning that survival was typically good in the apical half of the cochlea, i.e. for cochlear frequencies < 2.0 kHz, where there can be significant loss of outer hair cells. Of course, the 2.0 kHz region is roughly 15 mm from the basal tip of the cochlea, and these apical regions may be challenging to reach for drugs delivered at the round window.

Another overarching trend was that the supporting cells showed a radial gradient of resistance to degeneration that basically mirrored that of the hair cells, i.e. the Deiters were the most vulnerable, followed by the outer pillar cells and then the inner pillar cells. Given that the inner pillars are the Lgr5-expressing cells in the adult inner ear (Chai, Xia et al. 2011), and thus most likely to be responsive to the trans-differentiation signals, it is promising, in a regeneration context, that the inner pillars appear to be exceptionally resistant to degeneration in human cochleas.

Despite the difficulties inherent in classifying hearing loss etiologies in deceased humans, who often have complex medical histories involving several serious diseases, the present data suggest that patients with a history of noise damage have poorer supporting-cell survival, for a similar degree of threshold shift, than those with hearing loss from ototoxic drugs (Figs. 4B,C and 6C,D). Furthermore, the ototoxic cases were generally less complicated by strial degeneration or loss of neural elements (Fig. 6). Indeed, the enhanced neural survival in the ototoxic cases may be at least partially due to the enhanced supporting-cell survival (Suzuka and Schuknecht 1988) by virtue of the neurotrophins they release (Suzuka and Schuknecht 1988, Sugawara, Corfas et al. 2005, Zilberstein, Liberman et al. 2012). As discussed above, there can be excellent supporting-cell survival for several years after complete hair cells loss from ototoxic drugs (Leake and Hradek 1988, Sugawara, Corfas et al. 2005), and the present study included one ototoxic case who died shortly after the treatment with remarkably normal supporting-cell survival and architecture despite massive OHC loss. It is perhaps unfortunate, therefore, that patients with hearing loss from ototoxic drugs were excluded from the Frequency Therapeutics trials of hair cell regeneration in favor of those with a history of noise exposure (McLean, Hinton et al. 2021). The results of the trial might have been more promising if the inclusion criteria had been different.

Our analysis also suggests that for a given degree of high-frequency PTA, those with the most steeply sloping audiograms have poorer supporting-cell survival than those with flatter audiometric profiles. Some studies suggest that a flat audiometric profile could indicate more strial involvement in the generation of the threshold shift (Pauler, Schuknecht et al. 1988, Schuknecht and Gacek 1993, Dubno, Eckert et al. 2013), and the presence of strial pathology would reduce the utility of any hair cell regeneration. Regardless of its contribution to the audiometric pattern, strial pathology is common in aging ears (Wu, O’Malley et al. 2020), and better pre-mortem diagnostics for strial pathology are needed to further refine inclusion criteria for any future trials of regenerative therapies.

## Acknowledgements

Research supported by a grant from the National Institute on Deafness and other Communicative Disorders (P50 DC 015857) as well as by a sponsored research agreement from Frequency Therapeutics. The assistance of MengYu Zhu in slide digitization is gratefully acknowledged.

## References Cited

Bramhall, N. F., F. Shi, K. Arnold, K. Hochedlinger and A. S. Edge (2014). “Lgr5-positive supporting cells generate new hair cells in the postnatal cochlea.” Stem Cell Reports 2(3): 311–322.

Bredberg, G. (1968). “Cellular pattern and nerve supply of the human organ of Corti.” Acta Otolaryngol: Suppl 236:231+.

Chai, R., A. Xia, T. Wang, T. A. Jan, T. Hayashi, O. Bermingham-McDonogh and A. G. Cheng (2011). “Dynamic expression of Lgr5, a Wnt target gene, in the developing and mature mouse cochlea.” J Assoc Res Otolaryngol 12(4): 455–469.

Dallos, P. and D. Harris (1978). “Properties of auditory nerve responses in absence of outer hair cells.” J. Neurophysiol. 41(2): 365–383.

deTorres, A., R. T. Olszewski, I. A. Lopez, A. Ishiyama, F. H. Linthicum, Jr. and M. Hoa (2019). “Supporting cell survival after cochlear implant surgery.” Laryngoscope 129(1): E36–E40.

Dubno, J. R., M. A. Eckert, F. S. Lee, L. J. Matthews and R. A. Schmiedt (2013). “Classifying human audiometric phenotypes of age-related hearing loss from animal models.” J Assoc Res Otolaryngol 14(5): 687–701.

Forge, A. (1985). “Outer hair cell loss and supporting cell expansion following chronic gentamicin treatment.” Hear Res 19(2): 171–182.

Henry, K. S., S. Kale and M. G. Heinz (2016). “Distorted Tonotopic Coding of Temporal Envelope and Fine Structure with Noise-Induced Hearing Loss.” J Neurosci 36(7): 2227–2237.

Johnsson, L. G. and J. E. Hawkins, Jr. (1976). “Degeneration patterns in human ears exposed to noise.” Ann Otol Rhinol Laryngol 85(6 PT. 1): 725–739.

Johnsson, L. G., J. E. Hawkins, Jr., T. C. Kingsley, F. O. Black and G. J. Matz (1981). “Aminoglycoside-induced cochlear pathology in man.” Acta Otolaryngol Suppl 383: 1–19.

Kamakura, T. J. T. O’Malley and J. B. Nadol, Jr. (2018). “Preservation of Cells of the Organ of Corti and Innervating Dendritic Processes Following Cochlear Implantation in the Human: An Immunohistochemical Study.” Otol Neurotol 39(3): 284–293.

Leake, P. A. and G. T. Hradek (1988). “Cochlear pathology of long term neomycin induced deafness in cats.” Hear Res 33(1): 11–33.

Liberman, M. C. (1978). “Auditory-nerve response from cats raised in a low-noise chamber.” J Acoust Soc Am 63(2): 442–455.

Liberman, M. C. and N. Y. Kiang (1978). “Acoustic trauma in cats. Cochlear pathology and auditory-nerve activity.” Acta Otolaryngol Suppl 358: 1–63.

Linthicum, F. H., Jr. and J. N. Fayad (2009). “Spiral ganglion cell loss is unrelated to segmental cochlear sensory system degeneration in humans.” Otol Neurotol 30(3): 418–422.

Lobarinas, E., R. Salvi and D. Ding (2013). “Insensitivity of the audiogram to carboplatin induced inner hair cell loss in chinchillas.” Hearing research.

McGill, T. J. I. and H. F. Schuknecht (1976). “Human cochlear changes in noise induced hearing loss.” Laryngoscope LXXXVI(9): 1293–1302.

McLean, W. J., A. S. Hinton, J. T. J. Herby, A. N. Salt, J. J. Hartsock, S. Wilson, D. L. Lucchino, T. Lenarz, A. Warnecke, N. Prenzler, H. Schmitt, S. King, L. E. Jackson, J. Rosenbloom, G. Atiee, M. Bear, C. L. Runge, R. H. Gifford, S. D. Rauch, D. J. Lee, R. Langer, J. M. Karp, C. Loose and C. LeBel (2021). “Improved Speech Intelligibility in Subjects With Stable Sensorineural Hearing Loss Following Intratympanic Dosing of FX-322 in a Phase 1b Study.” Otol Neurotol 42(7): e849–e857.

McLean, W. J., X. Yin, L. Lu, D. R. Lenz, D. McLean, R. Langer, J. M. Karp and A. S. B. Edge (2017). “Clonal Expansion of Lgr5-Positive Cells from Mammalian Cochlea and High-Purity Generation of Sensory Hair Cells.” Cell Rep 18(8): 1917–1929.

Mellado Lagarde, M. M., G. Wan, L. Zhang, A. R. Gigliello, J. J. McInnis, Y. Zhang, D. Bergles, J. Zuo and G. Corfas (2014). “Spontaneous regeneration of cochlear supporting cells after neonatal ablation ensures hearing in the adult mouse.” Proc Natl Acad Sci U S A 111(47): 16919–16924.

Merchant, S. N., M. J. McKenna, J. C. Adams, J. B. Nadol, Jr., J. Fayad, R. Gellibolian, F. H. Linthicum, Jr., A. Ishiyama, I. Lopez, G. Ishiyama, R. Baloh and C. Platt (2008). “Human temporal bone consortium for research resource enhancement.” J Assoc Res Otolaryngol 9(1): 1–4.

Merchant, S. N. and J. B. Nadol (2010). Schuknecht’s Pathology of the Ear, 3rd Edition. Shelton, CT, People’s Medical Publishing House - USA.

Mizutari, K., M. Fujioka, M. Hosoya, N. Bramhall, H. J. Okano, H. Okano and A. S. Edge (2013). “Notch inhibition induces cochlear hair cell regeneration and recovery of hearing after acoustic trauma.” Neuron 77(1): 58–69.

Oesterle, E. C. and S. Campbell (2009). “Supporting cell characteristics in long-deafened aged mouse ears.” J Assoc Res Otolaryngol 10(4): 525–544.

Pauler, M., H. F. Schuknecht and J. A. White (1988). “Atrophy of the stria vascularis as a cause of sensorineural hearing loss.” Laryngoscope 98(7): 754–759.

Raphael, Y. and R. A. Altschuler (1991). “Scar formation after drug-induced cochlear insult.” Hear Res 51(2): 173–183.

Rask-Andersen, H., W. Liu and F. Linthicum (2010). “Ganglion cell and ‘dendrite’ populations in electric acoustic stimulation ears.” Adv Otorhinolaryngol 67: 14–27.

Resnik, J. and D. B. Polley (2021). “Cochlear neural degeneration disrupts hearing in background noise by increasing auditory cortex internal noise.” Neuron 109(6): 984–996 e984.

Schmiedt, R. A. (1989). “Spontaneous rates, thresholds and tuning of auditory-nerve fibers in the gerbil: comparisons to cat data.” Hear Res 42(1): 23–35.

Schuknecht, H. F. and M. R. Gacek (1993). “Cochlear pathology in presbycusis.” Ann Otol Rhinol Laryngol 102(1 Pt 2): 1–16.

Shera, C. A., C. Bergevin, R. Kalluri, M. M. Laughlin, P. Michelet, M. van der Heijden and P. X. Joris (2011). “Otoacoustic Estimates of Cochlear Tuning: Testing Predictions in Macaque.” AIP Conf Proc 1403: 286–292.

Slepecky, N. B., C. G. Henderson and S. Saha (1995). “Post-translational modifications of tubulin suggest that dynamic microtubules are present in sensory cells and stable microtubules are present in supporting cells of the mammalian cochlea.” Hear Res 91(1-2): 136–147.

Smith-Cortinez, N., R. Yadak, F. G. J. Hendriksen, E. Sanders, D. Ramekers, R. J. Stokroos, H. Versnel and L. V. Straatman (2021). “LGR5-Positive Supporting Cells Survive Ototoxic Trauma in the Adult Mouse Cochlea.” Front Mol Neurosci 14: 729625.

Sugawara, M., G. Corfas and M. C. Liberman (2005). “Influence of supporting cells on neuronal degeneration after hair cell loss.” J Assoc Res Otolaryngol 6(2): 136–147.

Suzuka, Y. and H. F. Schuknecht (1988). “Retrograde cochlear neuronal degeneration in human subjects.” Acta Otolaryngol Suppl 450: 1–20.

Taudy, M., Syka, Popelar and Ulehlova (1992). “Carboplatin and Cisplatin Ototoxicity in Guinea Pigs.” Audiology 31: 293–299.

Taylor, R. R., D. J. Jagger and A. Forge (2012). “Defining the cellular environment in the organ of Corti following extensive hair cell loss: a basis for future sensory cell replacement in the Cochlea.” PLoS One 7(1):p e30577.

Tsuji, J. and M. C. Liberman (1997). “Intracellular labeling of auditory nerve fibers in guinea pig: central and peripheral projections.” J Comp Neurol 381(2): 188–202.

Warchol, M. E. and J. T. Corwin (1996). “Regenerative proliferation in organ cultures of the avian cochlea: identification of the initial progenitors and determination of the latency of the proliferative response.” J Neurosci 16(17): 5466–5477.

Wu, P. Z., J. T. O’Malley, V. de Gruttola and M. C. Liberman (2020). “Age-Related Hearing Loss Is Dominated by Damage to Inner Ear Sensory Cells, Not the Cellular Battery That Powers Them.” J Neurosci 40(33): 6357–6366.

Wu, P. Z., W. P. Wen, J. T. O’Malley and M. C. Liberman (2019). “Assessing fractional hair cell survival in archival human temporal bones.” Laryngoscope.

Xu, S. A., R. K. Shepherd, Y. Chen and G. M. Clark (1993). “Profound hearing loss in the cat following the single co-administration of kanamycin and ethacrynic acid.” Hear Res 70(2): 205–215.

Zilberstein, Y., M. C. Liberman and G. Corfas (2012). “Inner hair cells are not required for survival of spiral ganglion neurons in the adult cochlea.” J Neurosci 32(2): 405–410.

